# Spatiotemporal dynamics in airborne fungi and fungal allergens across the United States

**DOI:** 10.64898/2025.12.23.696190

**Authors:** Sarah M. Gering, Ann M. Dillner, Scott Copeland, Noah Fierer

**Affiliations:** Department of Ecology and Evolutionary Biology, University of Colorado Boulder, Colorado, USA; Cooperative Institute for Research in Environmental Sciences, University of Colorado Boulder, Colorado, USA; Air Quality Research Center, University of California, Davis, Davis, CA 95616; Cooperative Institute for Research in the Atmosphere, Colorado State University, Ft. Collins, CO, USA

**Keywords:** aerobiology, fungal bioaerosols, aeroallergens, allergenic fungi, air quality monitoring networks

## Abstract

A broad diversity of fungi can be found in the near-surface atmosphere, with both the amounts and types of airborne fungi varying across space and time. However, the specific spatiotemporal patterns in airborne fungal assemblages often remain unquantified. This knowledge gap is particularly notable for allergenic fungi, despite the relevance of airborne fungal allergen exposures to public health. To better understand how airborne fungi, including known allergens, vary across broad spatial and temporal scales, we leveraged a pre-existing air sampling network to obtain bioaerosol samples from 7 national parks representing diverse biome types across the United States, with samples collected from each site every three days over an entire calendar year. We used marker gene DNA sequencing and quantitative PCR to characterize fungal assemblage composition and concentrations, including the concentrations of ten known allergenic genera. As expected, the composition of the airborne fungal assemblages, including the amounts and types of known allergens, varied across biomes. We also observed substantial temporal variation in the total amounts and types of fungi detected, particularly at higher latitude sites. While some of the temporal variation in allergen abundances followed seasonal patterns, we detected pronounced daily and weekly fluctuations in taxon-specific allergen abundances, with higher wind speeds generally associated with higher fungal allergen concentrations. Together, these results expand our understanding of fungal aerobiology across natural ecosystems and demonstrate how combining extensive air-sampling efforts with DNA-based analyses can improve assessments of public health risks associated with exposure to allergenic fungi in outdoor air.

**IMPORTANCE:** Despite major advances in sampling methodologies, sequencing technologies, and access to air quality monitoring networks, comprehensive assessments of the spatial and temporal variation in airborne fungi remain limited, even though fungi constitute a substantial proportion of the aerobiome. This is true even for allergenic fungal taxa, despite their importance to human health. Using DNA-based approaches, we characterized fungal assemblages in the near-surface atmosphere over a single calendar year at 7 national parks across the US. We found that each site had distinct airborne fungal assemblages with unique temporal patterns in fungal abundances and composition. Substantial spatiotemporal variation was also observed for known fungal allergens, driven by both seasonal trends and environmental factors. This study advances our understanding of the ecological patterns that structure airborne fungal communities and the factors influencing outdoor exposures to allergenic fungi, improving our ability to assess and predict health risks associated with fungal allergens.

## INTRODUCTION

Fungi are ubiquitous and abundant in the near-surface atmosphere as many fungal spores can become airborne and be transported over distances ranging from meters to thousands of kilometers (Anees-Hill et al., 2022; Chaudhary et al., 2022). These airborne fungal spores are most commonly sourced from fungi living on leaf surfaces (Levetin & Dorsey, 2006) or in soils (Bowers et al., 2009; Fröhlich-Nowoisky et al., 2009), with the dissemination of fungal spores through the atmosphere a key mode of fungal dispersal, including the dispersal of fungi that can have direct impacts on ecosystem and human health (Brown & Hovmøller, 2002). Most notably, many fungal allergens are transported through the atmosphere, where they can trigger allergies and asthma in susceptible individuals. Such allergic diseases are estimated to impact the health and well-being of 25% of the US population (Ng & Boersma, 2023) with a similarly high prevalence of allergic diseases in other regions across the globe (Shin et al., 2023). There is a long history of research on airborne fungi (Martinez-Bracero et al., 2022), including studies focused on allergens (Al-Shaarani & Pecoraro, 2024), given their ecological importance and their relevance to public health. By combining extensive air sampling efforts with DNA sequencing-based analyses, it is now feasible to comprehensively quantify spatiotemporal patterns in airborne fungal diversity, including allergenic fungi, at broader scales and with more consistent, higher resolution taxonomic assignments than possible with more traditional approaches (Abrego et al., 2024; Tordoni et al., 2021; Tournayre et al., 2025; Yamamoto et al., 2014).

Total airborne fungal concentrations and concentrations of individual taxa found in the near-surface atmosphere can vary strongly as a function of geographic location (Bowers et al., 2013; Chaudhary et al., 2022). This spatial variation can be apparent at a range of scales and is most commonly associated with numerous factors that differ across locations including: climate and vegetation type (Marčiulynas et al., 2023), soil conditions (Pan et al., 2025), and land-use types or practices (Lin et al., 2018) with these factors collectively affecting the distributions of fungal taxa in source environments, the release of fungal spores into the air, and the likelihood of fungal spores persisting in outdoor air (Segers et al., 2023). While these and other measured or unmeasured variables can contribute to spatial variation in airborne fungal concentrations, we often lack the comprehensive data on taxon-specific spatial patterns in fungal concentrations needed to build a predictive understanding of how and why airborne fungal diversity may vary across geographic locations. These knowledge gaps are even more apparent for airborne fungal allergens. As many allergy-sufferers are acutely aware, different locations harbor different amounts of outdoor allergens, but, in contrast to our extensive understanding of allergenic pollen distributions (Ren et al., 2022) our understanding of how concentrations of specific fungal allergenic taxa vary across regions remains limited.

Even at a given location, the amounts and types of fungi found in outdoor air can vary appreciably over time. This temporal variation may be apparent across a range of time scales - from diurnal patterns (Fierer et al., 2008) to seasonal patterns (Nicolaisen et al., 2017) depending on the taxon and the specific site characteristics. For example, fungal allergen abundances often exhibit pronounced seasonality, contributing to seasonal variation in the prevalence of allergic symptoms (Priyamvada et al., 2017), with the timing and magnitude of the temporal variation depending on the location and taxa in question (Fukutomi & Taniguchi, 2015; Priyamvada et al., 2017). Several factors can contribute to this temporal variation including the timing of spore release (Norros et al., 2023; Savage et al., 2012), phenological patterns in fungal growth in the source environments (Krah et al., 2023), and meteorological conditions such as temperature, rainfall, humidity, and snow cover (Liu et al., 2020; Lu et al., 2022; Núñez et al., 2021; Tamiya et al., 2025). These factors can collectively contribute to observed changes in airborne fungal assemblages over time in ways that are often difficult to predict *a priori* (Green et al., 2006; Grewling et al., 2019).

Here, we used air filters (PM_10_) collected by a pre-existing air quality monitoring network (IMPROVE; Malm et al., 1994, https://vista.cira.colostate.edu/Improve/improve-program/) to investigate spatial and temporal patterns in total airborne fungal concentrations and diversity, with a focus on allergenic fungi given their public health importance. A total of 836 filter samples were included in this study with these samples collected every three days for an entire year (2021) from each of 7 national parks across the US, which span a broad range of ecosystem types that are minimally affected by human impacts (Fig. 1). These samples were analyzed using cultivation-independent DNA-based approaches (quantitative PCR and ITS marker gene sequencing) to answer three questions. First, how do airborne fungal assemblages vary across different biomes and how do the observed temporal patterns within sites vary throughout the year? Second, how do the concentrations of known allergenic fungi vary across sites and within and between seasons at individual sites? Third, what environmental factors, including meteorological conditions, are most closely associated with observed temporal variation in allergenic fungal concentrations at individual sites? Our study builds on a growing body of research to expand our understanding of fungal aerobiology and the occurrence of allergenic fungal taxa in the near-surface atmosphere.

**Fig. 1:**
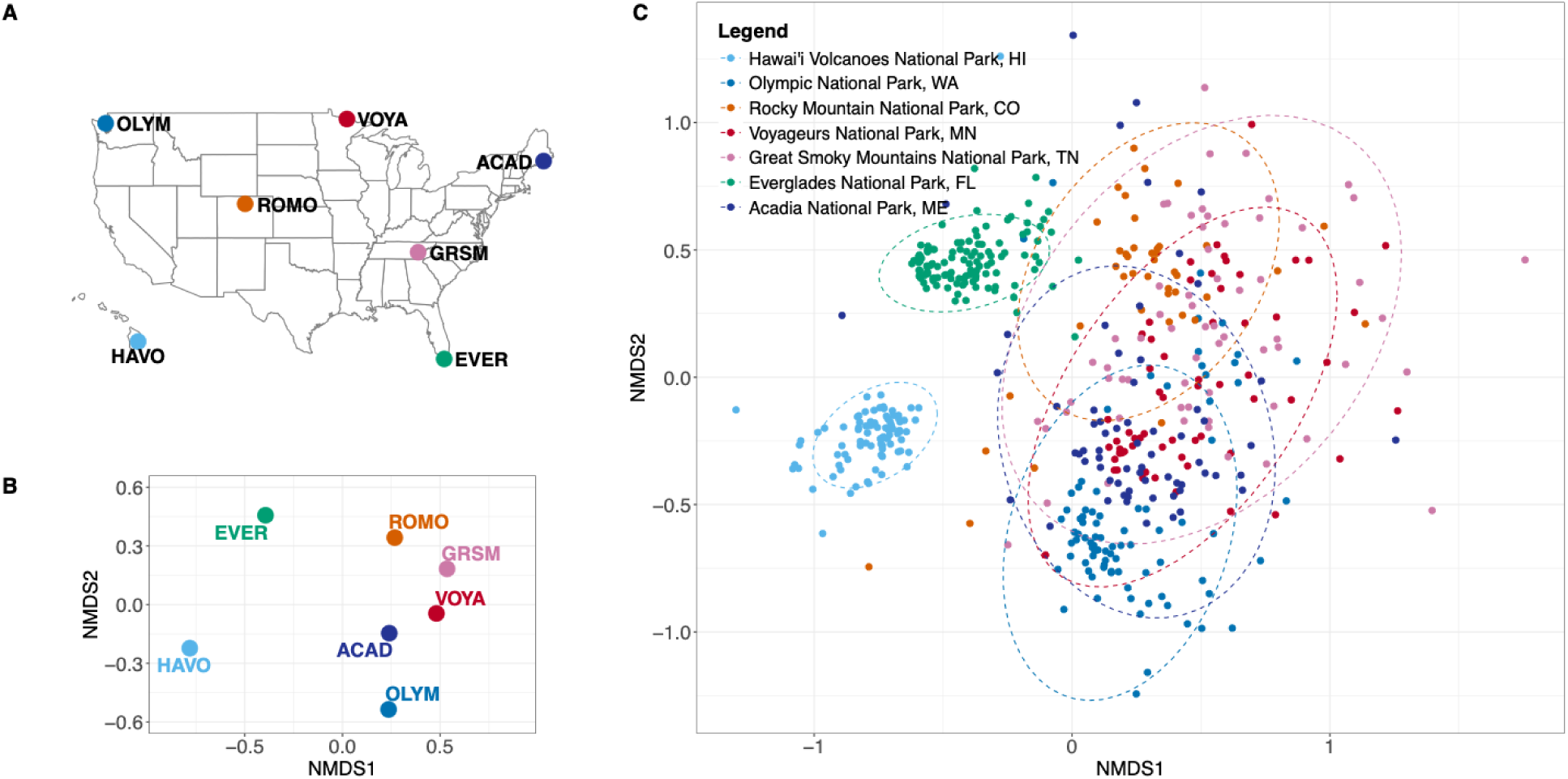
Map of the sampling locations and fungal community composition across sites. (A) Map of the United States showing all 7 sampling sites. Sites are labeled as acronyms and colored consistently throughout the study include Hawai’i Volcanoes National Park, Hawaii (HAVO, light blue); Olympic National Park, Washington (OLYM, ocean blue); Rocky Mountain National Park, Colorado (ROMO, orange); Voyageurs National Park, Minnesota (VOYA, red); Great Smoky Mountains National Park, Tennessee (GRSM, pink); Everglades National Park, Florida (EVER, green); and Acadia National Park, Maine (ACAD, dark blue). (B) Nonmetric multidimensional scaling (NMDS) ordination of the centroid of each site and (C) NMDS ordination of all samples colored by site. The dashed ellipses represent a 95% confidence showing the distribution of the community composition at each national park.

## RESULTS AND DISCUSSION

### Site and Sample Set Description

In this study, we quantified the variation observed in the fungal aerobiome found in the near-surface atmosphere across 7 locations (Fig. 1A) over a calendar year. By examining dissimilarity in fungal assemblage composition across sites over time (Fig. 1B, 1C), we expanded our understanding of the seasonal and environmental patterns that shape airborne fungi, including known allergenic taxa. Of the 836 air filter samples collected for this project, 514 samples met our threshold for inclusion in downstream analyses based on qPCR and sequence data (see Methods). These 7 sites were selected because they are located in distinct biomes with minimal direct human impacts. All sites and reported results are referred throughout as the following: HAVO (Hawai’i Volcanoes National Park), OLYM (Olympic National Park), ROMO (Rocky Mountain National Park), VOYA (Voyageurs National Park), GRSM (Great Smoky Mountains National Park), EVER (Everglades National Park), and ACAD (Acadia National Park) (Fig. 1A, Supplementary Table 1).

### Methodological considerations

Before detailing our results, it is important to highlight some important caveats. First, our sampling and analytical approaches do not allow us to distinguish between airborne spores and fragments of fungal tissues that could contribute to the airborne DNA pool, nor could we differentiate between viable and non-viable fungi in the collected air samples. Second, some of the fungal DNA captured on the filters may have degraded during the 24-h sampling period or during extended storage of the filters prior to DNA extraction. Although all samples were stored under identical conditions and with near-identical timing across sites, samples from individual sites varied in the time they were archived (12-24 months), with those collected at the end of 2021 spending less time in storage than those collected in early 2021. Thus, we wanted to determine if differential storage times may have affected our ability to resolve temporal variation within sites (Fig. S2). Seasonal patterns in total fungal DNA concentrations (Fig. 2), observed fungal richness (Fig. S1), and assemblage dissimilarity (Fig. 1C) were stronger than any apparent effect of storage duration. While we cannot rule out the possibility of storage effects, such effects are unlikely to limit our ability to assess temporal patterns.

**Fig. 2:**
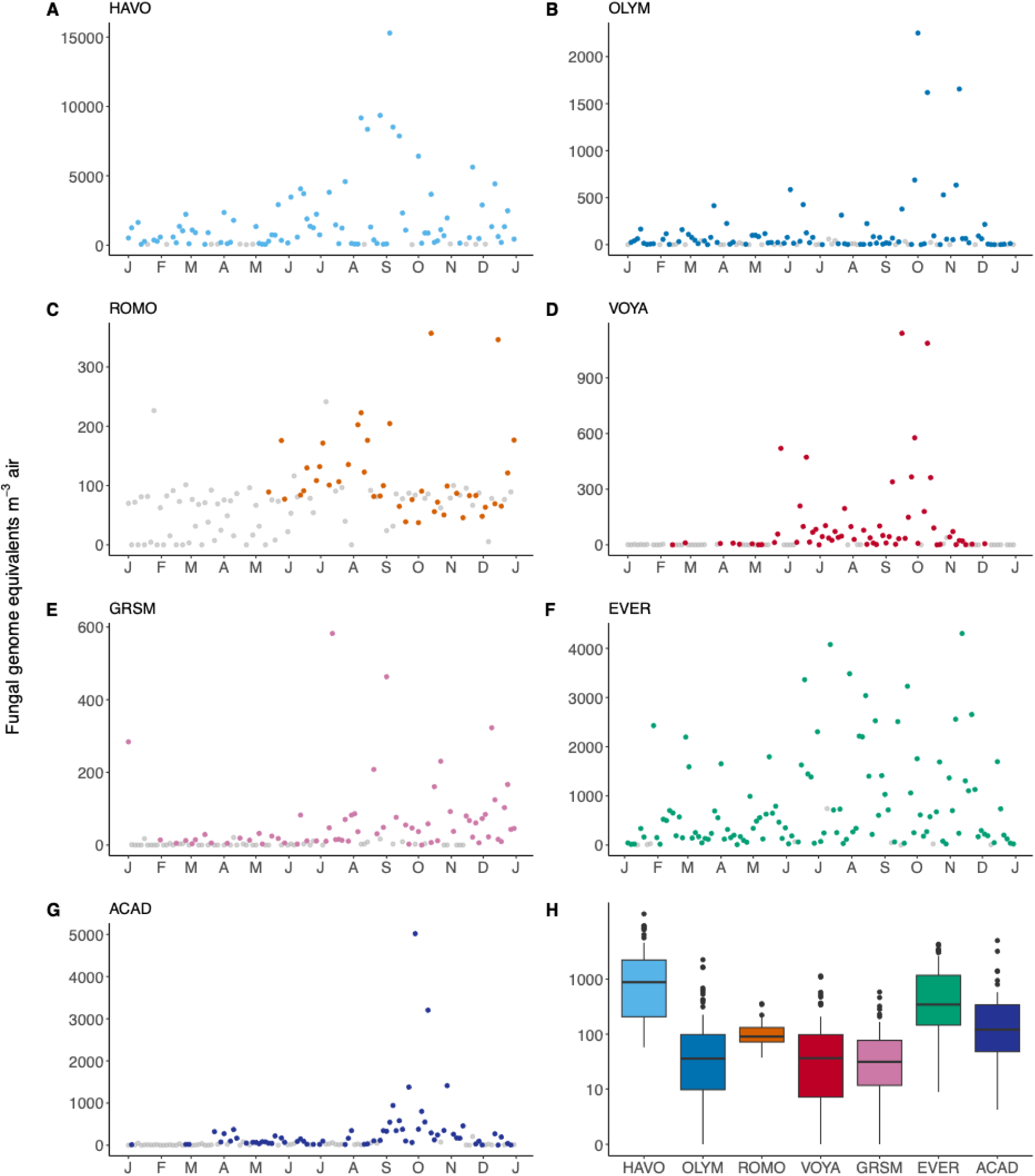
Quantitative PCR-based measurements of total fungal DNA concentrations over time across all 7 sites. Panels (A-G) include temporal qPCR patterns, with the grey points depicting samples below detection for sequencing. Colored points represent samples with successful qPCR amplification and sequencing that were retained for downstream analyses. Panel (H) displays pseudo-log-scaled boxplots summarizing the median total fungal DNA concentrations across sites calculated from samples that were used in downstream analysis (in color).

We also note that sample coverage varied across sites. More than 70% of the samples from the time series at EVER, HAVO, and OLYM were included in downstream analyses, while ∼55% were retained from VOYA, GRSM, and ACAD, and only 34% of samples from ROMO met our criteria for inclusion (see Methods). In particular, many of the samples collected during the winter months from ROMO, VOYA and ACAD did not yield sufficient fungal DNA for qPCR or sequencing (Fig. 2C, 2D, 2G). This pattern is expected as the sites would often be snow covered during these time periods which should greatly reduce fungal spore release from plants and soil. More generally, our inability to collect complete time series data from all sites reflects some of the challenges inherent in conducting aerobiological studies, especially where fungal concentrations can be very low during winter months (Tamiya et al., 2025; Wu et al., 2025).

### Spatial patterns in total fungal abundances and assemblage composition

We first focused on geographic differences in fungal assemblages across each of the 7 sampling locations. On average, HAVO and EVER, the tropical and subtropical sites, had higher average fungal concentrations (Fig. 2A and 2F) than the more temperate sites (Fig. 2B-E; 2G, 2H). These general patterns align with previous studies, showing that more tropical sites tend to have elevated total fungal concentrations in the near-surface atmosphere (Abrego et al., 2024). The overall composition of the airborne fungal assemblages also differed between sites with location explaining ∼25% of overall variation (PermANOVA, R^2^ = 0.25, p < 0.001, Fig. 1B). While we observed considerable temporal variation within sites (discussed below), the airborne fungal assemblages at HAVO and EVER were consistently distinct from those found at the other sites (HAVO: pairwise PermANOVA R^2^ = 0.27-0.30, p<0.001, EVER: pairwise PermANOVA R^2^ = 0.11-0.28, p<0.001) (Fig. 1C). For those sites with elevated temporal variation, including VOYA, GRSM and ACAD, the overall site-specific differences in assemblage composition were weaker, but still significant (R² = 0.04-0.06, p<0.001) as evident from Fig. 1B and 1C. These geographic differences in airborne fungal assemblages are not surprising given that these sites represent such a broad range of biome types with distinct soils, vegetation types, and climates – all factors that are known to contribute to biogeographic patterns in airborne fungi (Fröhlich-Nowoisky et al., 2012; Nicolaisen et al., 2017; Schmale & Ross, 2015).

The geographic patterns in fungal assemblages are also evident from differences in the relative abundances of the fungal classes across sites (Fig. 3). A few fungal classes were typically dominant at all sites, including members of the Agaricomycetes, Pezizomycetes, and Dothideomycetes classes (Fig. 3), results that align with previous studies (Woo et al., 2018). However, the relative abundances of these taxa generally exhibited site-specific differences despite the observed intra-site temporal variation. For example, members of the class Agaricomycetes were typically the most abundant at nearly all sites, representing 70% of reads on average, especially at HAVO or OLYM where they represented 88% of reads on average (Fig. 3A and 3B). Pezizomycetes was the second most abundant class across all sites ranging from an average of 1% at HAVO up to 35% of reads at ROMO (Fig. 3). At finer levels of taxonomic resolution, the site-specific differences in fungal assemblages are even more evident. For example, <0.5% of fungal ASVs (20 out of 15890 ASVs in total) were detected in all 7 sites with these 20 ASVs spanning a range of genera including *Pyronema*, *Cladosporium*, *Aspergillus*, *Epicoccum*, and *Alternaria* (Table S2). Likewise, only 7% (1156) of the 15890 ASVs were detected across >3 sites. Together these results highlight the high degree of variation in airborne fungal assemblages across biomes, variation that is consistent with results from previous studies (Krah et al., 2023; Tipton et al., 2022; Woo et al., 2018).

**Fig. 3:**
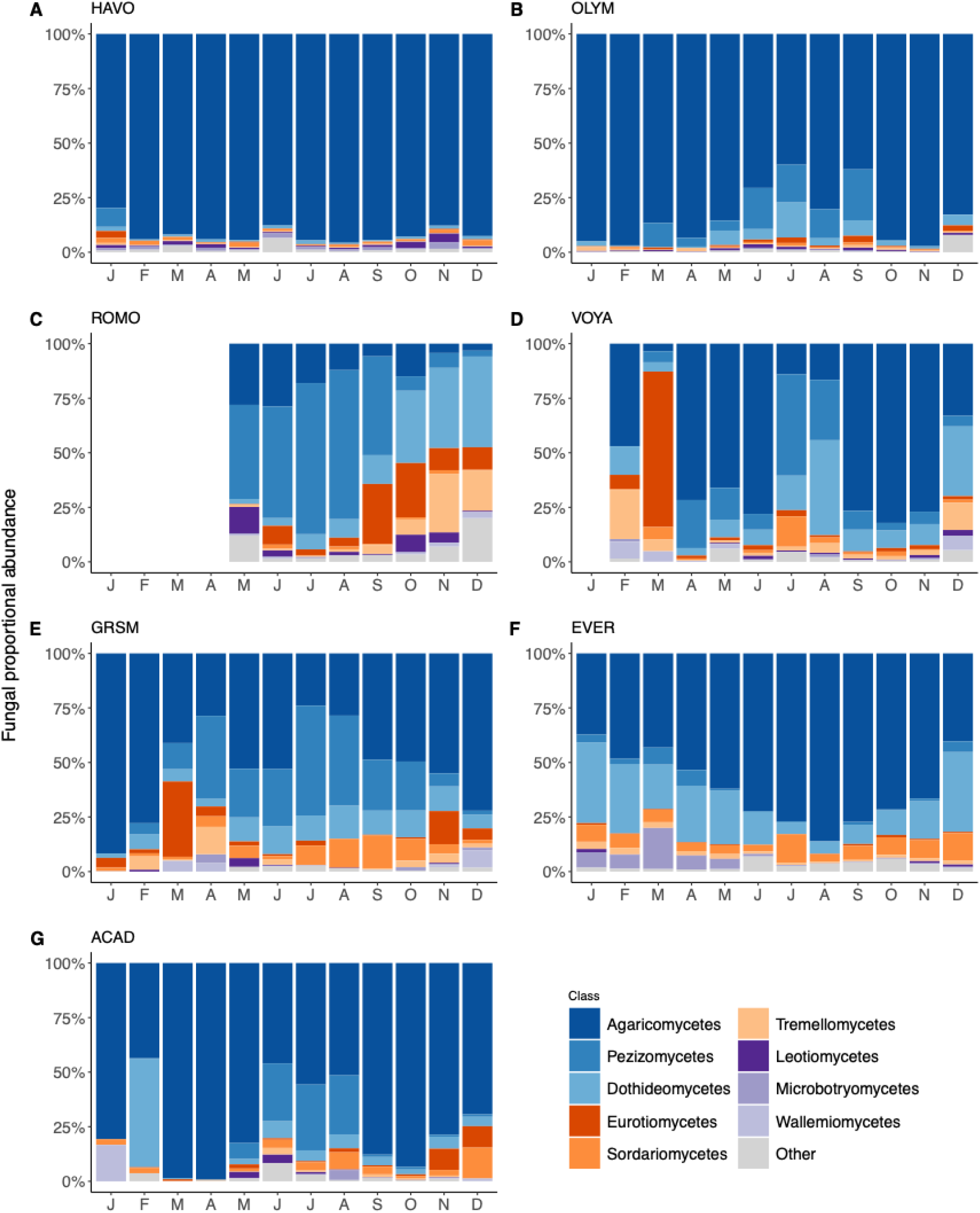
Mean proportional abundances of fungal classes averaged by month across all 7 sites for the full year. Panels (A-G) are labeled by the site from left to right: Hawaii Volcanoes National Park, Hawaii (HAVO), Olympic National Park, WA (OLYM), Rocky Mountain National Park (ROMO), Acadia National Park, Maine (ACAD), Everglades National Park, FL (EVER), Voyageurs National Park, MN (VOYA) and Great Smoky Mountains National Park, TN (GRSM). ROMO and VOYA do not have bars during winter months because too many samples were excluded during quality filtering, likely due to low fungal biomass during winter conditions.

### Temporal variation in airborne fungi

While we focused on the broad site-level differences in airborne fungi above, there is clearly substantial temporal variation at individual sites in total fungal concentrations (Fig. 2), observed richness (Fig. S1) and overall fungal assemblage composition (Fig. 1, Fig. 3). The temporal variation in assemblage composition was low at HAVO (Fig. 1C, 3A), as expected given the relatively stable climate at this tropical site (Kodama et al., 2024), but seasonal patterns were evident at the other sites with previous studies noting similar seasonality (Abrego et al., 2024; Nageen et al., 2023). Fungal DNA concentrations and observed richness generally peaked during the late summer and fall months at many of the sites (HAVO, OLYM, ROMO, VOYA, and ACAD) (Fig. 2, Fig. S2), a pattern that is likely related to a combination of warmer soil temperatures and the timing of leaf fall (Ortega-Rosas et al., 2025). We also observed that several sites showed later summer increases in Pezizomycetes relative to Agaricomycetes (OLYM, GRSM, ACAD; Fig. 3B, 3E, 3G), contrary to studies that have reported that Pezizomycetes tend to become more abundant during cooler months (Kumari et al., 2016). Nearly all sites exhibited seasonal patterns, although the specific timing of the seasonal patterns varied depending on the site, likely related to differences in climate and the phenology of spore release (Magyar et al., 2016). Even within seasons, we observed pronounced day-to-day variation in the amounts and types of fungi detected in the near-surface atmosphere (Fig. 2). This daily and weekly variation emphasizes the importance of collecting high-resolution time series when studying the fungal aerobiome and emphasizes that airborne fungi are sensitive to changes in environmental conditions, including atmospheric conditions, that are likely a product of taxon-specific phenological patterns, differences in the rates at which fungal spores are aerosolized from their respective source environments, and differences in atmospheric residence times that are difficult to predict *a priori* (Lagomarsino et al., 2020).

### Spatiotemporal variation in allergenic fungi

We next examined the spatiotemporal patterns in the amounts and types of airborne allergenic fungi given their public health importance. These analyses focused on the 10 known allergenic fungal genera that were most common across our samples (Esch et al., 2001; Fig. 4A-4G and Fig. S3A-S3G). These 10 genera collectively represented between 1.3% of the total fungi at HAVO up to 21.5% of total fungi at ROMO (Fig. 4H), with daily values ranging from 0% up to 99% of all fungi detected depending on the site and time of year (Supplementary Fig. 3A-3G). The most abundant allergenic fungi across all sites were members of the *Cladosporium*, *Alternaria*, and *Aspergillus* genera, taxa that have been shown to be dominant in other fungal aerobiome surveys (Anees-Hill et al., 2022). However, there was appreciable site-level variation in the abundances of particular allergenic genera (Fig. 4H). For example, *Botrytis* was the dominant allergenic genus at HAVO and EVER had higher abundances of *Curvularia* and *Fusarium* than the other sites. These geographic differences in taxon-specific allergen concentrations are likely a product of differences in local source environments (including vegetation type) and climate conditions (Choi et al., 2023; Olsen et al., 2019). Likewise, total allergen abundances (summed across 10 allergenic genera) varied appreciably across sites with the subtropical site (EVER) that is relatively warm and humid having allergen abundances that were at up to five times higher than all the other sites (Fig. 4H).

**Fig. 4:**
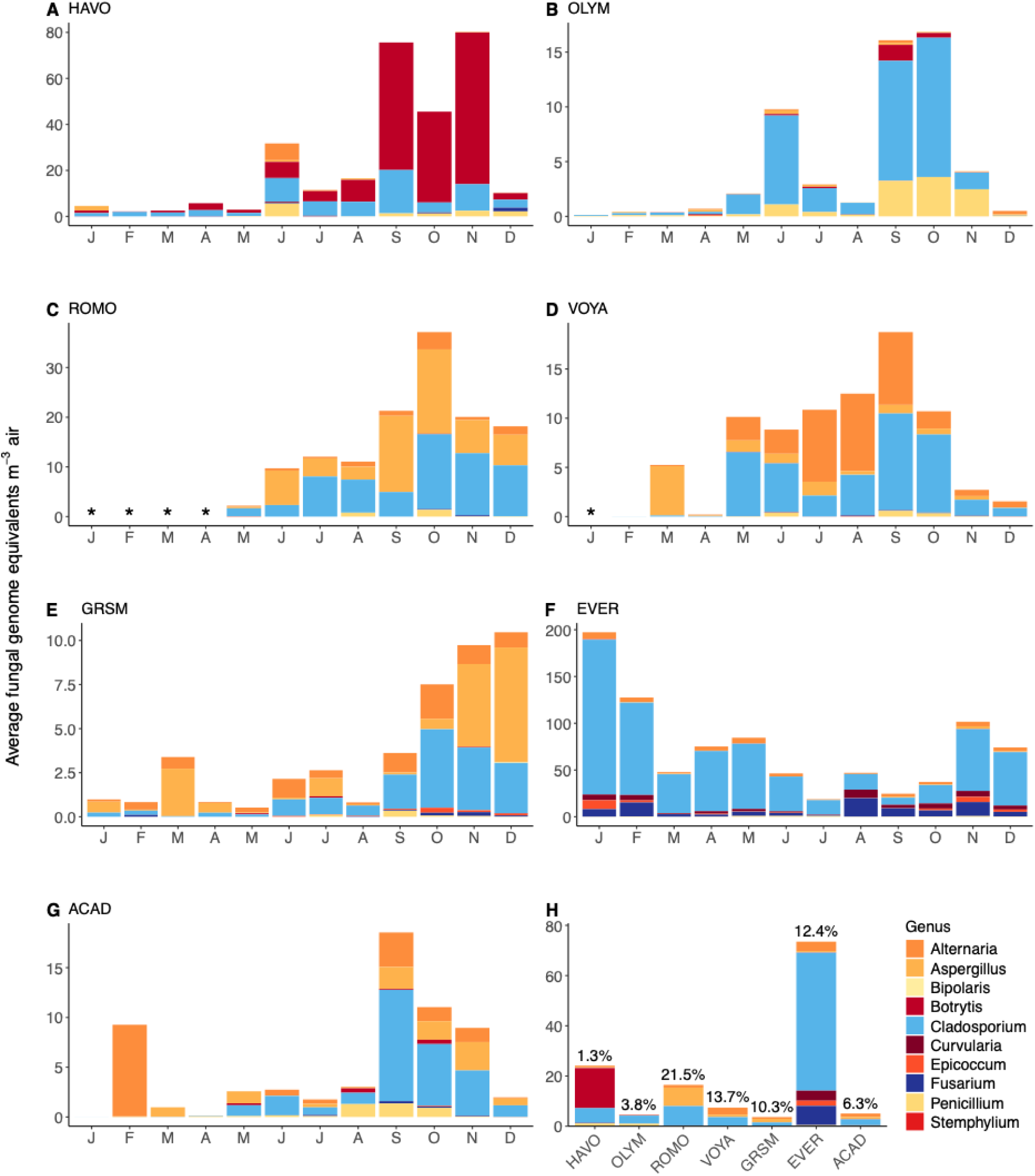
Average monthly DNA amounts represented by allergenic fungal genera. Panels (A-G) include the average concentrations of allergenic fungal genera across all 7 sites by month. Panel (H) displays mean concentrations of allergenic-associated fungal genera across sites as stacked bars by genus with percentages above each site indicating the percent abundance of the ten allergenic fungi relative to all other fungi. Asterisks indicate months where allergenic taxa were absent or below detection limits.

Although the dominant allergens were relatively consistent within sites over time (Fig. 4 and Fig. S3), there was appreciable temporal variation in total allergen concentrations. Some of this temporal variation was seasonal, but the degree of seasonality varied depending on the site in question. More specifically, we show that the sampling month (our proxy for season), was significantly associated with temporal variation in total allergen abundances across 6 of the 7 sites (Table 1). The monthly variation was strongest at ACAD (R^2^ = 0.39), intermediate at HAVO, OLYM, ROMO, VOYA, and GRSM (R^2^ = 0.09-0.15, p<0.05), and weak and non-significant at EVER (Table 1). Total allergen abundances at most sites (except for EVER) peaked in late summer through fall months (August through November) but the specific timing and duration of this peak varied across sites (Fig. 4). Our observation that most sites have the highest airborne allergen concentrations in late summer and fall months is consistent with our overall fungal patterns (Fig. 2) as well as results reported in previous work (Anees-Hill et al., 2022; Grewling et al., 2019; Symon et al., 2025). Further, although fungi are not the sole contributor to allergies, we note that the peak in fungal allergen concentrations in late summer and fall months corresponds to increases in the prevalence of seasonal allergies during a similar time period across many US metropolitan areas (Stallard-Olivera & Fierer, 2024).

**Table 1:** Environmental factors tested that explain airborne fungal allergen abundances across the 7 national park sites. Seasonality R² values represent variance explained by month using generalized additive modeling. The environmental variables are listed with their correlation direction (+ positive, - negative), individual R² values, and p-values. ‘Full Model R²’ represents total variance when combining seasonality and environmental variables. A p-value <0.05 is considered a significant result.

| Site | Seasonality R<sup>2</sup> | Environmental variables | + or - | R<sup>2</sup> | p-value | Full Model R<sup>2</sup> |
| --- | --- | --- | --- | --- | --- | --- |
| HAVO | 0.126 | Nitrate (NO<sub>3</sub>) | + | 0.087 | 0.008 | 0.321 |
|  |  | Precipitation | - | 0.066 | 0.029 |  |
|  |  | Wind Gusts | + | 0.060 | 0.022 |  |
| OLYM | 0.133 | Sodium (Na) | + | 0.068 | 0.024 | 0.202 |
| ROMO | 0.136 | Soil moisture | - | 0.261 | 0.001 | 0.228 |
|  |  | Evapotranspiration | - | 0.218 | 0.003 |  |
|  |  | Wind Gusts | + | 0.124 | 0.030 |  |
|  |  | Sodium (Na) | - | 0.123 | 0.031 |  |
| VOYA | 0.092 | Wind Gusts | + | 0.186 | 0.009 | 0.400 |
|  |  | Minimum temperature | + | 0.121 | 0.008 |  |
|  |  | Dust Metric | + | 0.096 | 0.020 |  |
|  |  | Evapotranspiration | + | 0.078 | 0.037 |  |
| GRSM | 0.146 | Humidity | - | 0.074 | 0.040 | 0.260 |
| EVER | 0.030 | Wind Gusts | + | 0.103 | 0.0006 | 0.172 |
|  |  | Sodium (Na) | + | 0.094 | 0.001 |  |
|  |  | Wind Speed | + | 0.073 | 0.004 |  |
|  |  | PM10 | + | 0.041 | 0.033 |  |
| ACAD | 0.386 | None | n/a | n/a | n/a | 0.386 |

### Environmental drivers of airborne fungal allergen concentrations

Although we observed pronounced seasonal patterns in total allergen abundances across most sites, month alone explained a relatively small amount of the observed temporal variation per site (Table 1) as there was considerable variation observed at weekly and daily time scales (Fig. S3). Thus, we next sought to determine what environmental variables might best explain the additional variance in total allergen abundances beyond sampling month (residual variance). We found that wind speed was often positively correlated with total allergen abundances. More specifically, maximum wind gusts and average wind speed were positively correlated with total allergen abundances at 4 of the 7 sites: HAVO, ROMO, VOYA, and EVER (R² = 0.06-0.19, p<0.05, Table 1). The association between wind speed and elevated concentrations of fungal allergens has been observed previously (Pellitier et al., 2025) and reflects the importance of wind to the aerosolization and dispersal of fungal spores (Cáliz et al., 2018; Schmale & Ross, 2015). Other environmental variables that were also observed to be associated with the residual variance in total allergen concentrations beyond sampling month included sodium concentrations which were positively correlated with allergen abundances at the coastal sites OLYM (R² = 0.068, p, <0.05) and EVER (R² = 0.094, p <0.001), likely reflecting shared dispersal mechanisms where windy conditions that facilitate fungal dispersal also transport marine aerosols (Alsante et al., 2021). We also identified negative associations with soil moisture at ROMO and with humidity at GRSM, results that align with the observation that release of allergenic fungal spores can be elevated under drier conditions (Priyamvada et al., 2017). Overall, seasonality and wind speed emerged as the most consistent predictors of the temporal variation in allergenic fungal concentrations across most of the sites, while other variables considered (including PM_10_ and PM_2.5_ concentrations) provided minimal explanatory power (Table 1). However, there were substantial amounts of temporal variation that could not be explained by our models, perhaps due to site-specific phenology patterns or local atmospheric dynamics that influence the aerosolization and residence time of allergenic spores in the near-surface atmosphere. More generally, our time-series analyses of fungal allergen abundances further highlight the complexity of fungal aerobiology and a need for more consistent and comprehensive monitoring of fungal allergen exposures (Williams & Smith, 2021).

## FUTURE DIRECTIONS

Our year-long sampling campaign conducted across 7 US national parks revealed extensive spatial and temporal variation in airborne fungal communities. Our findings add to a growing body of research characterizing fungal aerobiology in natural ecosystems, with site-specific differences depending on location and time of year. Most of the locations studied exhibited seasonal patterns, with airborne fungal concentrations peaking from late summer to fall, and seasonality and wind emerging as the primary drivers of elevated allergenic fungal concentrations. Future research would benefit from linking airborne fungal allergen concentrations to public health data to better understand when allergies might be most prevalent and, ultimately, build a predictive understanding of the spatiotemporal dynamics in fungal-triggered allergy symptoms.

## MATERIALS AND METHODS

### Sample Collection

Air filter samples were collected by the Interagency Monitoring of Protected Visual Environments (IMPROVE) network from 7 sites (Fig. 1A, Table S1) managed by the US National Park Service. The selected sites are situated in distinct biomes and span a wide range in climates and vegetation types (Table S1).

The air filters used in this study were collected between January 01, 2021 and December 31, 2021, using polytetrafluoroethylene (PTFE) filters (3 μm pore size, 25 mm diameter; Measurement Technologies Laboratory) fitted to a sampling apparatus with a PM_10_ inlet, which is positioned ∼4 meters above the ground. Air was pulled through each filter at a rate of 17 L min^-1^ for 24 hours, with samples taken every third day (IMPROVE, 2022). Total filtered air volumes averaging 24.2 m^3^ day^-1^ (range 20.3 to 25.7 m^3^ day^-1^). The PM_10_ air filters collected by the IMPROVE network are used solely for gravimetric mass measurements of particulate matter (PM) and are collected concurrently with PM_2.5_ filters that are used for chemical analyses (IMPROVE, 2020). The filters were archived at room temperature between 12 to 24 months and, once shipped, were stored at −20°C until DNA was extracted. All samples analyzed in this study were stored under identical conditions but were assessed for potential effects of storage duration (see above and Fig. S2). IMPROVE provided the aerosol chemical data associated with each sample with relevant chemical metrics used in this study, including PM_10_ and PM_2.5_ mass, organic carbon, sodium (Na), sulfate (SO_4_), nitrate (NO_3_), potassium (K), and a composite mineral of fine soil as a dust metric concentration (Malm et al., 1994). Chemical data were accessed and downloaded from https://views.cira.colostate.edu/fed/default.aspx. For all 7 sites, we downloaded daily meteorological data from Visual Crossing Weather (Visual Crossing, 2024) for the year 2021. The meteorological data used for our analyses included minimum and maximum daily temperatures, average daily temperature, relative humidity, precipitation, average wind speed, maximum wind speed, and average wind direction during each sampling period. We also included soil moisture levels and evapotranspiration metrics for each sampling period for all sites (NASA, 2025).

### DNA Extractions

We extracted DNA from 114-122 filters per site from each of the 7 sites for a total of 836 filters plus associated field blanks using the ZymoBIOMICS 96 DNA Kit. Filters were cut into quarters using flame-sterilized scissors, aseptically transferred to the ZR BashingBead™ Lysis Tubes, 650 μL of the Bashing Bead Buffer was added, and then the tubes were vortexed on a horizontal vortexer for 40 min at maximum speed. The tubes were centrifuged at 10,000 x g for 1 min, and between 400-500 μL supernatant was transferred into the 96-well block, where 1200 μL of the ZymoBIOMICS DNA Binding Buffer was added. The 96-well block was sealed with a sterile foil cover and vortexed upright on a horizontal vortexer for 2 minutes. The plate sat at room temperature for 10 min to ensure binding before the next transfer step. We followed the manufacturer’s instructions for the rest of the procedure, except that, at the final step, we added 50 μL of ZymoBIOMICS DNase/RNase Free water, omitted the step with the Silicon HRC Plate, then re-eluted the eluate onto the column matrix. We then incubated the plates for three minutes and centrifuged again per the manufacturer’s recommendations to maximize DNA recovery. Five to six extraction blanks were randomly assigned into wells on each extraction plate to check for potential contaminants introduced during the extraction process, following ‘best practices’ for working with low-biomass samples (Fierer et al., 2025).

### ITS amplicon sequencing

To characterize the fungal assemblages associated with each filter, we PCR-amplified the extracted DNA using barcoded primers targeting the fungal internal transcribed spacer region (ITS1; Emerson et al., 2015). Our PCR protocol followed that previously described (Gering et al., 2024). PCRs were prepared in duplicate 25 μL reactions consisting of 12.5 μL Platinum™ II Hot-Start PCR Master Mix (Invitrogen, Carlsbad, CA, USA), 7.5 μL PCR H_2_O, 1 μL of the 10 μM barcoded primer, and 4 μL of template DNA. PCR cycling conditions were at 94°C for 2 min, followed by 35 cycles of 94°C for 15 sec, 60°C for 15 sec and 68°C for 1 min, with a final extension step at 72°C for 10 min. Samples were then cleaned and normalized using the SequalPrep normalization kit (Thermo Fisher Scientific, Carlsbad, CA, USA) and pooled in equimolar concentrations. Pooled libraries were sequenced on two separate Illumina MiSeq runs (Illumina, California, USA) using the 2×250bp cycle kit at the Center for Microbial Exploration at the University of Colorado Boulder. In total, we extracted and sequenced 836 air samples, plus 19 field blanks, 60 DNA extraction blanks and 11 PCR blanks. Raw reads were processed via the DADA2 pipeline (v. 4.1.; Callahan et al., 2016), merged, quality-filtered, and trimmed to 240 bp. Amplicon sequence variants (ASVs) were determined as reads that shared 100% sequence identity (v. 138.1; Quast et al., 2012; Yilmaz et al., 2014) and all ASVs were classified against the UNITE fungal taxonomy reference database (v. 10.05.2021; Nilsson et al., 2019).

Any samples, including negative controls, that contained <3000 reads were removed prior to downstream analyses, along with ASVs represented by <10 reads in total across all samples and any ASVs not classified to the phylum level of resolution. Out of the 836 air filters collected across all 7 sites that were extracted and sequenced (in addition to 60 extraction blanks, 19 field blanks, and 11 no template controls), we retained 514 air samples that met our quality control thresholds, which resulted in a final sample size of n = 85 (HAVO), n = 85 (OLYM), n = 41 (ROMO), n = 61 (VOYA), n = 64 (GRSM), n = 112 (EVER), and n = 66 (ACAD), one field blank, no PCR blanks, and six extraction blanks. Across the retained air filter samples, there were a total of 9,755,471 ITS reads, representing 15,890 unique ASVs, with a median of 17,150 quality-filtered reads per air sample. Seven negative control samples (out of the 90 total blanks/controls processed) passed through this quality filtering, with these 7 ‘blank’ samples having a median of 4,754 reads per blank. We looked for potential external contaminants and only removed a single ASV, classified as a member of the *Wallemia* genus, from the full dataset as it was detected in at least three of the blank samples, but was observed in low abundance in the air samples (100 reads or 1.51% of reads per sample on average). All samples were rarefied to a minimum read depth of 3009 for downstream analyses.

### Quantitative PCR

To estimate the total amount of fungal DNA on each filter sample, we used quantitative PCR (qPCR) on a Bio-Rad CFX Connect real-time system (Bio-Rad Laboratories, Hercules, CA, USA) using the same ITS1-targeting fungal primers described above. Following previous lab methods tested on IMPROVE air filters (Gering et al., 2024), the reactions were performed in 25 μL reactions containing 12.5 μL 2X master mix (Thermo Scientific SYBR Green), 1.25 μL of forward and 1.25 μL of reverse primers, 6 μL of PCR-grade water, and 4 μL of DNA. We included two no-template controls on each 96-well plate and generated standard curves using genomic DNA from *Aspergillus fumigatus*. Fungal qPCR thermocycling conditions were as follows: 95°C for 15 min, with 40 cycles of denaturation at 94°C for 45 sec, annealing at 55°C for 1 min, and extension at 72°C for 1:30 min, followed by a final extension step at 72°C for 10 min. All qPCR results are reported as *A. fumigatus* genome equivalents m⁻³ air.

Samples underwent both qPCR and amplicon sequencing. Only those samples that passed our quality control thresholds (≥3000 sequencing reads and quantifiable qPCR results) were included in downstream community analyses. We note that the samples that did not meet both thresholds and were therefore excluded from downstream analyses are depicted as grey points in Fig. 2A-2G to illustrate both technical limitations and seasonal patterns in sample coverage across our dataset. The summary averages and reported values for qPCR were determined from only the retained samples, as indicated by the colored points (Fig. 2A-G). It is important to note that qPCR cannot account for gene copy number variation in mixed fungal communities (Hospodsky et al., 2010). Therefore, our results serve as a proxy for biomass to identify overall spatial and temporal patterns, rather than being interpreted as true absolute fungal (or spore) abundances in the air.

### Allergen Analyses and Approach

We focused our allergen analyses on ten known respiratory allergens, including the following genera: *Aspergillus, Bipolaris, Botrytis, Cladosporium, Curvularia, Epicoccum, Fusarium, Penicillium,* and *Stemphylium (*Esch et al., 2001). This list does not encompass all known fungal allergens, but does capture those commonly considered important triggers of allergic responses (Kwong et al., 2023). However, we note that not all members of these genera are necessarily allergenic, and we are measuring DNA concentrations associated with allergenic fungal genera, not concentrations of specific fungal antigens (or spores). To estimate fungal concentrations (via qPCR) of these 10 allergenic genera, we first calculated their relative abundance from the sequence data by summing the genus-level ASV reads and then dividing by the total number of fungal reads per sample. We then multiplied the taxon-specific relative abundances with the qPCR estimates of total fungal DNA concentrations to quantify the absolute abundances of allergenic genera (reported in fungal genome equivalents m^-3^ of air per sample). We used rarefied sequence data for these analyses, but rarefied and non-rarefied results were well-correlated (Pearson correlation, R^2^ = 0.99).

### Statistical Analyses

All analyses were conducted in R (v. 2024.12.1.563, R Core Team, 2025). The R packages used included *phyloseq* (v. 1.52.0, McMurdie & Holmes, 2013), *tidyr* (v. 1.3.1, Wickham et al., 2019) and *dplyr* (v. 1.1.4, Wickham et al., 2023) for data organization, *vegan* (v. 2.7-1, Oksanen et al., 2023) and *mgcv* (v. 1.9-3; Wood, 2025) for GAM modeling and *ggplot2* (v. 3.5.2, Wickham, 2016), *cowplot* (v. 1.2.0, Wilke, 2025), *ggpubr* (v. 0.6.2, Kassambara, 2023), and *gridextra* (v. 2.3, Auguie & Antonov, 2017) for visualizations. We used permutational multivariate analysis of variance (PermANOVA) with Bray-Curtis dissimilarity, using the ‘adonis2’ function in *vegan* to compare the taxonomic composition of the fungal assemblages across sites. To identify potential environmental and seasonal variables associated with fungal allergen abundance data, we used generalized additive models (GAMs, Wood, 2025). The allergen abundance data were log-transformed with a +1 pseudocount, and seasonal patterns were assessed using the cyclic cubic splines approach by month, per the *mgcv* package. We cross-compared different dimensions of k with the Akaike Information Criterion (AIC) to determine an appropriate smoothing dimension, identifying a k = 8, which improved our model fit.

We assessed potential storage effects on community composition over time by averaging Bray-Curtis dissimilarities between all samples and the average of the oldest ten samples at each site to assess whether storage time had an impact on community composition over time (Fig. S2). We visualized these trends in dissimilarity using locally weighted scatterplot smoothing (LOESS).

To identify which environmental and atmospheric variables might account for additional temporal variation in allergen abundances beyond season (i.e. month-to-month variation), we extracted the residuals from the seasonal generalized additive model (GAM) results at each site and correlated these residuals with our environmental variables using regression analysis. For a complete list of all the variables tested, see Supplementary Table S3, as this list includes both weather variables (e.g. air temperature, humidity, wind direction/speed) as well as general aerosol chemical measurements such as PM_2.5_ and PM_10_ associated with each sample. Additionally, we incorporated other atmospheric chemical variables into our GAM models to determine what chemical characteristics might be directly or indirectly associated with the observed temporal variation in allergen abundances.

## ACKNOWLEDGMENTS

We thank all IMPROVE-affiliated researchers and field scientists associated with sample collection and processing. IMPROVE is a collaborative association of state, tribal, and federal agencies, and international partners. The US Environmental Protection Agency is the primary funding source, with contracting and research support from the National Park Service. The Air Quality Group at the University of California, Davis is the central analytical laboratory, with ion analysis provided by the Research Triangle Institute and carbon analysis provided by Desert Research Institute. We gratefully acknowledge support from the NSF Biology Integration Institutes Program under Award # 2120117. We thank the CSU One Health Institute for contributions to and support of Bii: Regional Onehealth Aerobiome Discovery Network (BROADN) research. We also thank the Cooperative Institute for Research in Environmental Sciences at the University of Colorado Boulder and the National Science Foundation Graduate Research Fellowship.

## COMPETING STATEMENT

The authors declare no competing financial interests.

## DATA AVAILABILITY

All sequence data generated for this project is in the process of being deposited in the Genbank Short Read Archive (BioProject SUB15874634).

## AUTHOR CONTRIBUTIONS

Sarah M. Gering, Conceptualization, Data curation, Formal analysis, Writing – original draft, Writing – review and editing | Ann M. Dillner, Data curation, Writing – review and editing | Scott Copeland, Data curation, Writing – review and editing | Noah Fierer, Conceptualization, Data curation, Formal analysis, Funding acquisition, Project administration, Supervision, Writing – review and editing

## SUPPLEMENTAL MATERIAL

**Supplementary Table 1.** Site summary details for all 7 sampling locations.

| Site | Latitude | Longitude | Elevation (m) | State | National Park (NP) | Biome |
| --- | --- | --- | --- | --- | --- | --- |
| HAVO | 19.4309 | -155.2579 | 1258 | Hawaii | Hawai'i Volcanoes NP | tropical rainforest |
| OLYM | 48.0065 | -122.9727 | 599 | Washington | Olympic NP | Pacific coast and temperate forest |
| ROMO | 40.2783 | -105.5457 | 2760 | Colorado | Rocky Mountain NP | montane |
| VOYA | 48.4132 | -92.8303 | 425 | Minnesota | Voyageurs NP | boreal forest and temperate deciduous forest |
| GRSM | 35.6334 | -83.9416 | 810 | Tennessee | Great Smoky Mountains NP | temperate deciduous forest |
| EVER | 25.3910 | -80.6806 | 1.25 | Florida | Everglades NP | subtropical wetlands |
| ACAD | 44.3800 | -68.2600 | 157 | Maine | Acadia NP | Atlantic coast, boreal and deciduous forest |

**Supplementary Fig. 1:**
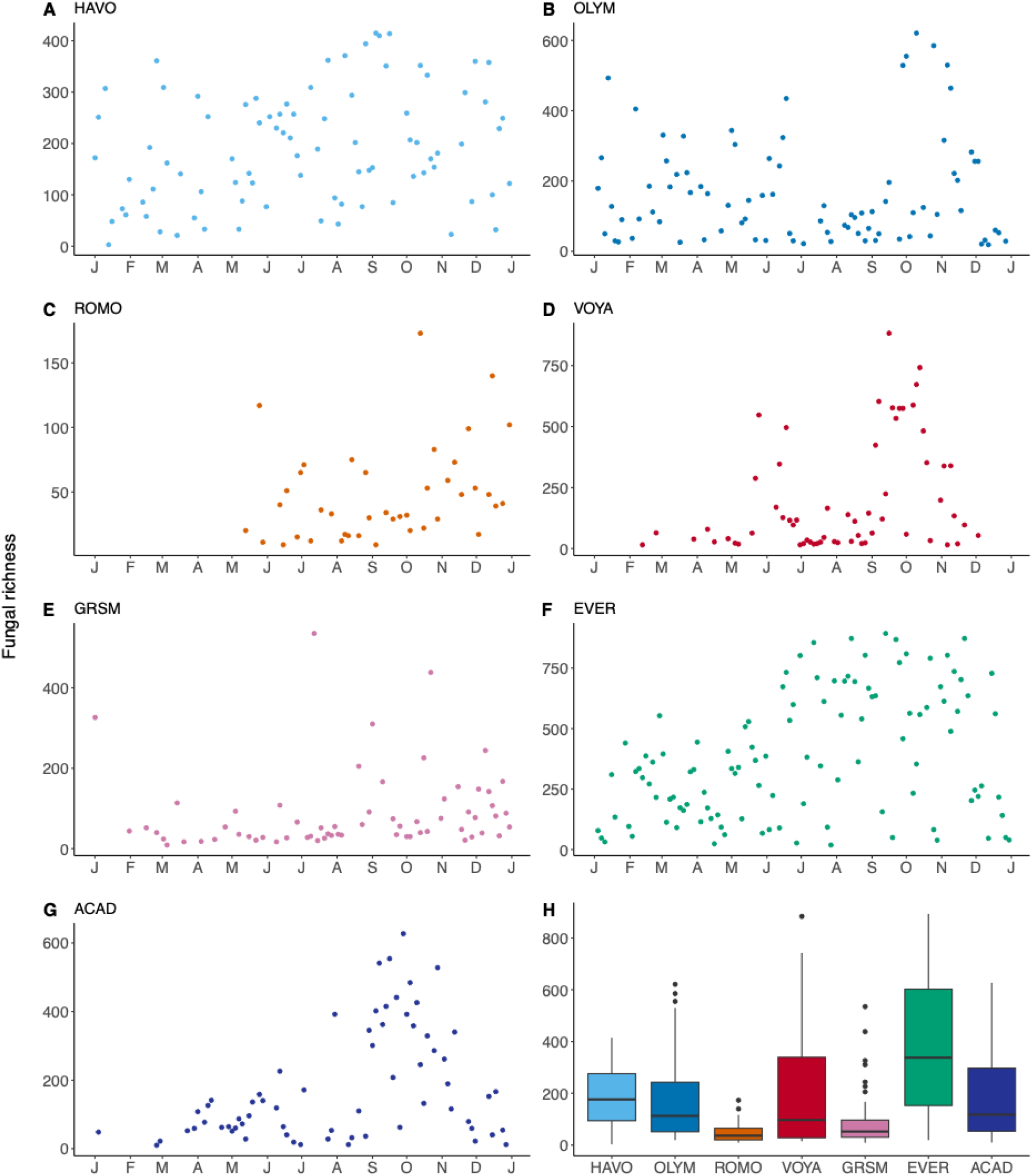
Fungal richness, measured as the number of observed taxa, over time across all 7 sites. Panels (A-G) show temporal patterns per site and panel (H) includes the median overall fungal richness by site.

**Supplementary Fig. 2:**
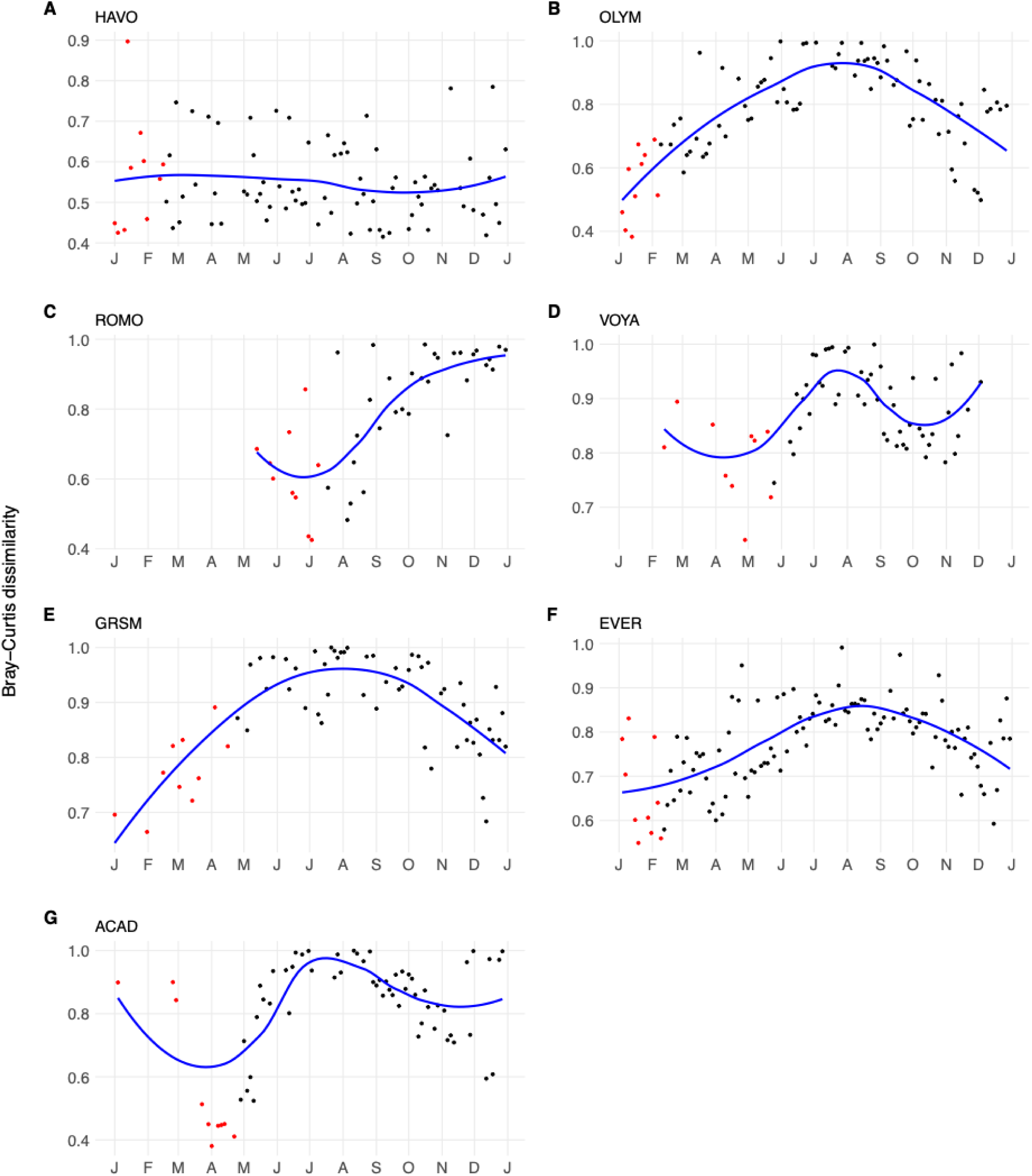
Bray-Curtis dissimilarity comparing all samples to the baseline (average of the oldest ten samples sequenced, shown in red) at each site to assess storage impact on archived filters. If storage time systematically altered community composition, we would expect dissimilarity to increase linearly over time as samples spent progressively less time in storage. Instead, dissimilarity patterns follow seasonal cycles at all sites, with values ranging from approximately 0.4 to 1.0 throughout the year. Samples at HAVO and EVER, contain the most complete annual sequencing coverage, demonstrate seasonal cycling rather than storage degradation. At ROMO and VOYA, early-year samples were not retained or sparse, so baseline samples were averaged during late spring/early summer with the linear increase in dissimilarity for later samples reflecting seasonal differences rather than storage artifacts and are shown here for transparency.

**Supplementary Fig. 3:**
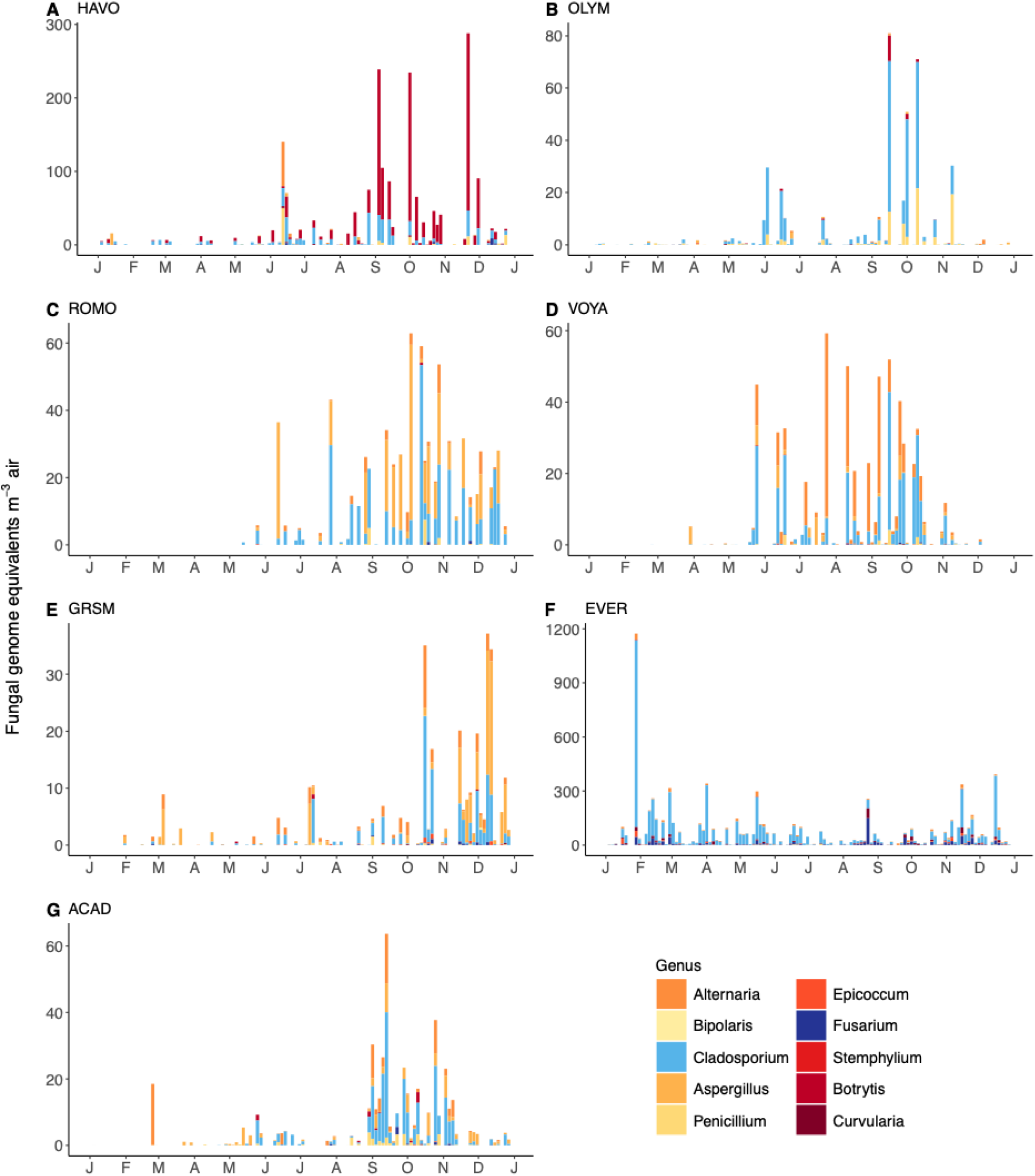
Total allergen abundances for each sampling day across sites in fungal genome equivalents m^3^ of air. Panels (A-G) are labeled by the site from left to right: Hawai’i Volcanoes National Park, HI (HAVO), Olympic National Park, WA (OLYM), Rocky Mountain National Park, CO (ROMO), Voyageurs National Park, MN (VOYA), Great Smoky Mountains National Park, TN (GRSM), Everglades National Park, FL (EVER), and Acadia National Park, ME (ACAD). Each stacked bar represents an individual sampling date, showing the DNA concentrations of allergenic fungal genera. Colors represent different allergenic genera as shown in the legend.

**Supplementary Table 2.**
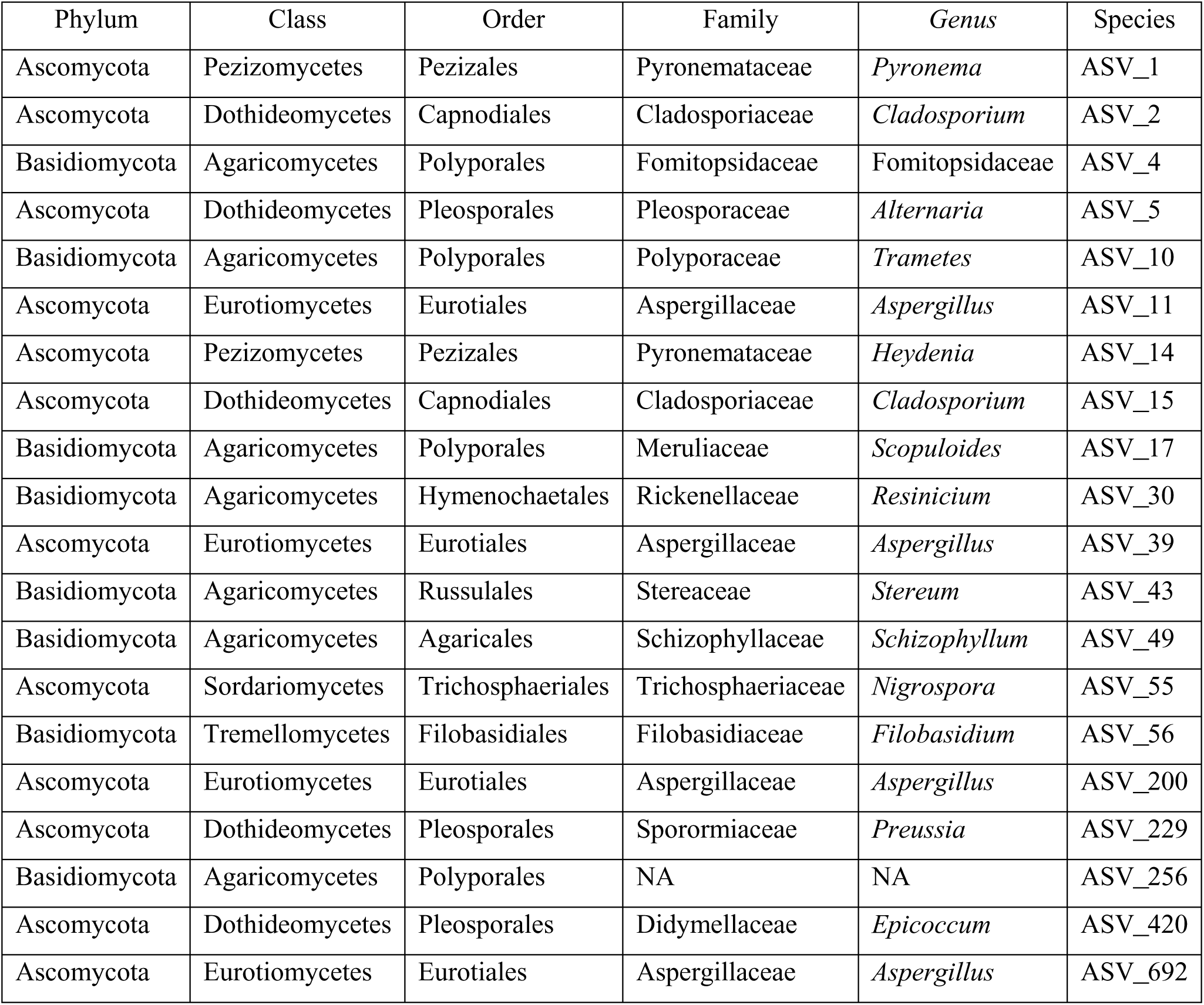
List of ASVs shared across all sites.

**Supplementary Table 3.** Complete list of explanatory variables tested, including weather variables, atmospheric chemistry measurements, and environmental variables associated with each sample.

| Weather Variables | Atmospheric Chemistry | Environmental Variables |
| --- | --- | --- |
| Air Temperature | PM<sub>2.5</sub> | Solar radiation |
| Minimum Temperature | PM<sub>10</sub> | Evapotranspiration |
| Maximum Temperature | SO<sub>4</sub> | Surface soil moisture |
| Relative Humidity | Na |  |
| Precipitation | K |  |
| Wind direction | K |  |
| Wind speed | NO<sub>3</sub> |  |
| Wind gusts | OC |  |
|  | S |  |
|  | Dust Composite metric* |  |

\*Dust composite metric includes aluminum (Al), silicon (Si), calcium (Ca), iron (Fe), and titanium (Ti).

